# Intrinsic Homeostatic Plasticity in Mouse and Human Sensory Neurons

**DOI:** 10.1101/2023.06.13.544829

**Authors:** Lisa A. McIlvried, John Smith Del Rosario, Melanie Y. Pullen, Andi Wangzhou, Tayler D. Sheahan, Andrew J. Shepherd, Richard A. Slivicki, John A. Lemen, Theodore J. Price, Bryan A. Copits, Robert W. Gereau

## Abstract

In response to changes in activity induced by environmental cues, neurons in the central nervous system undergo homeostatic plasticity to sustain overall network function during abrupt changes in synaptic strengths. Homeostatic plasticity involves changes in synaptic scaling and regulation of intrinsic excitability. Increases in spontaneous firing and excitability of sensory neurons are evident in some forms of chronic pain in animal models and human patients. However, whether mechanisms of homeostatic plasticity are engaged in sensory neurons under normal conditions or altered after chronic pain is unknown. Here, we showed that sustained depolarization induced by 30mM KCl induces a compensatory decrease in the excitability in mouse and human sensory neurons. Moreover, voltage-gated sodium currents are robustly reduced in mouse sensory neurons contributing to the overall decrease in neuronal excitability. Decreased efficacy of these homeostatic mechanisms could potentially contribute to the development of the pathophysiology of chronic pain.

## INTRODUCTION

In order to maintain stable long-term function, neuronal networks must adapt in the face of mounting experience-dependent plasticity. Two main mechanisms of homeostatic plasticity have been proposed as master regulators of neuronal activity: synaptic scaling and homeostatic regulation of intrinsic excitability ^1, 2^. Synaptic scaling, which has been reported in neocortical, hippocampal, and spinal cord neurons ^1, 3, 4^, refers to an overall adjustment of synaptic strength that serves to compensate for perturbations in neuronal network activity. These adjustments occur in all synapses on a neuron, scaling up or down endogenous ligand-gated receptors, thus normalizing overall neuronal firing. Alternatively, homeostatic regulation of intrinsic excitability is a non-synaptic homeostatic mechanism that controls individual cell activity by dynamically modulating intrinsic neuronal excitability, generally through changes in ion channel expression or function^2, 5, 6^. These adaptations have been observed in excitatory-induced neuronal cells from human pluripotent stem cells, neuronal cells from cultured hippocampal and cortical mouse neurons, rodent myenteric neurons that innervate regions of the enteric nervous system, tectal neurons from the *Xenopus* retinotectal circuit, and pyloric neurons from the pyloric circuit in the crab, *Cancer borealis*^2, 7–13^.

In animal models of chronic pain, changes in neuronal excitability are observed throughout the pain neuraxis, including peripheral nociceptive fibers, neurons of the dorsal horn of the spinal cord, neurons in the central nucleus of the amygdala, and pyramidal neurons in the anterior cingulate cortex^14–22^. In addition, human sensory neurons from patients experiencing neuropathic pain also become hyperexcitable, suggesting that adaptive alterations in intrinsic neuronal excitability are present in humans^23–25^. Thus, in both rodent and human, sensory neurons from individuals with persistent pain show increased firing rates and changes in intrinsic excitability. This sustained increase in activity begs the question of whether sensory neurons engage homeostatic mechanisms to regulate intrinsic excitability.

In the present study, we investigated whether mechanisms of homeostatic plasticity can be engaged in mouse and human dorsal root ganglia (DRG) sensory neurons. We found that sustained depolarization (24h, 30mM KCl) leads to adaptive alterations in neuronal excitability in mouse and human sensory neurons. The inhibition of intrinsic excitability includes changes in rheobase, input resistance, and action potential (AP) waveform. These changes were reversible within 24 hours - a necessary requirement of a homeostatic plasticity process. We also found that sustained depolarization produced only a modest alteration in voltage-gated potassium currents, but a robust reduction in voltage-gated sodium currents in mouse sensory neurons. This finding suggests that voltage-gated sodium currents contribute to the decrease in neuronal excitability and potentially serve as a regulatory mechanism to drive homeostatic control in sensory neurons. Understanding the intrinsic molecular mechanisms that contribute to this homeostatic downregulation of peripheral sensory neuron excitability could aid in understanding the role of these processes in restoring normal sensation in the context of injury, and whether disruption of such mechanisms might contribute to the development of chronic pain.

## RESULTS

### Sustained depolarization of mouse DRG neurons decreased neuronal excitability

To understand if homeostatic plasticity is engaged in sensory neurons, cultured DRG neurons at 4 days *in vitro* (DIV) were exposed to 30mM KCl for 24h to induce sustained neuronal depolarization^26–30^ (Fig. 1A, **top**). This treatment had no impact on the viability of the cultured DRG neurons (Fig. 1A, **bottom left**), and resulted in sustained depolarization of the cells to −15 ± 0.5mV (n=5) without driving activity (Fig. 1A, **bottom right**).

**Figure 1:**
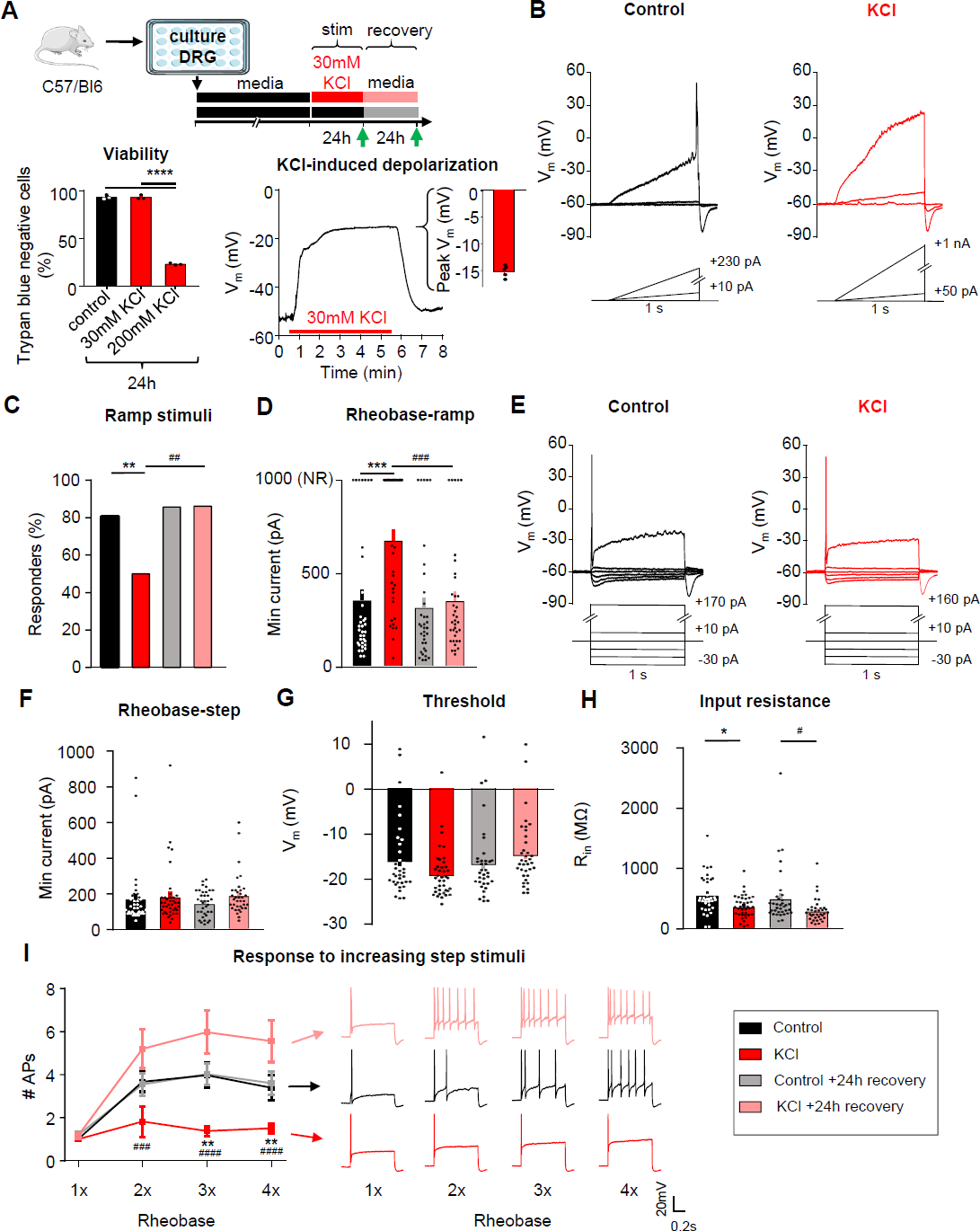
Sustained depolarization of mouse DRG neurons decreased neuronal excitability. A) *Top*: Experimental design for testing homeostatic regulation of intrinsic excitability to a sustained depolarization stimulus in cultured, small diameter DRG neurons from C57Bl/6 mice. *Bottom left*: Viability of mouse DRG cultures after 24hr stimulation with control, 30mM or 200mM KCl treatments (N=3 mice; Trypan Blue staining performed in replicate for each culture condition). *Bottom right*: example current-clamp trace from acute, 5min application of 30mM KCl to a mouse DRG neurons and subsequent washout of effect. KCl application resulted in a sustained depolarization to - 15 ± 0.5mV (n=5 neurons). B) Example current clamp traces from a control cultured mouse sensory neuron (left; black) or one treated for 24hrs with media containing 30mM KCl (right; red). Current injections were delivered in ramp pulses, indicated below traces, in 10pA increases until a reliable action potential was achieved (responders) or reaching the maximum of 1nA without firing (non-responders; NR). Comparison of the proportion of responders (C) and rheobase from all cells (D) following treatment. E) Example current clamp traces from the same cultured small-diameter mouse DRG neurons as in A, with current injections delivered in step pulses. All cells responded. Comparison of rheobase (F), threshold (G), input resistance (H), and number of APs fired in response to 1-4 times rheobase current injections with example traces (I), determined from step pulses. Treatment groups: 24h in control, media alone (n=36; black), 24h in media supplemented with 30mM KCl (n=39; red), 24h control followed by an additional 24h in fresh media (n=35; grey), and 24h 30mM KCl followed by an additional 24h “recovery” in fresh media (n=36; pink). *=significant difference between control and KCl groups; ^#^=significant comparison between KCl and KCl+24h recovery groups; one-way and two-way ANOVAs with Tukey’s multiple comparisons. */^#^ p<0.05, **/^##^ p<0.01, ***/^###^ p<0.001, ****/^####^ p<0.0001. Data are represented as mean ± SEM

Excitability was assessed using whole-cell patch-clamp electrophysiology after 24 ± 4 hrs of incubation in 30mM KCl in culture media, or after a further 24 ± 4hrs recovery in fresh media (Fig. 1A, **green arrows**^31^). The recovery group allows us to assess for a reversal in excitability to the sustained stimulus, which is considered a hallmark indication of homeostatic plasticity. Control neurons received media changes only. Data were collected from 146 small diameter (20.6± 0.02 μm) DRG neurons (<30 μm, putative nociceptors^32–35^) from 8 mice (4 male, 4 female). There were no significant differences in excitability between male and female mice, therefore data were pooled.

In current-clamp recordings (Fig. 1B), the majority of neurons in control media fired an action potential (AP) to a ramp depolarization (29/36 or 81%), and only half of the neurons fired APs to this stimulus after 24h incubation in 30mM KCl (19/38 or 50%; Fig. 1C). This effect recovered within 24h (31/36 or 86% responder rate) when neurons were placed back in fresh control media. KCl-treated DRG neurons had a significantly higher rheobase compared to the media control group, and this effect also recovered within 24h in fresh control media (Fig. 1D). The neurons that did not fire any AP up to the maximum injected current ramp of 1nA were classified as non-responders (**NR;** Fig. 1B**-right and 1D-top**) and rheobase was set to 1nA (Fig. 1D**-top of the graph**). We also assessed rheobase, AP threshold, and response to suprathreshold stimuli (accommodation) to 1-second step current injections (Fig. 1E). We found that there were no significant changes in rheobase (Fig. 1F) or in AP threshold (Fig. 1G), but there was a significant change in input resistance (Fig. 1H) using the step protocol, possibly reflecting an increase in channel insertion into the membrane. In addition, there was a significant decrease in the number of action potentials generated at three and four times rheobase in KCl-treated DRG neurons compared to DRG neurons treated with control media. This effect recovered within 24h in fresh control media (Fig. 1I). The increase in rheobase and the decrease in the percentage of DRG neurons responding to the ramp current-injection protocol as well as the decrease in input resistance and to suprathreshold step stimuli in the KCl-treated group show that there is an overall decrease in neuronal excitability of sensory neurons following a sustained depolarization. The reversibility of these effects following 24h washout of the depolarizing stimulus suggests that somatosensory neurons can engage a dynamic form of homeostatic plasticity.

The passive and active electrophysiological properties of the cells were also examined. There were no changes in resting membrane potential (V_rest_) or whole-cell capacitance (pF) between KCl and control-treated groups (Table 1). Analysis of AP characteristics showed that KCl-treated DRG neurons lost their shoulder, which is a distinctive inflection that occurs during the descending phase of the action potential (Fig. 2 A-C) ^36, 37^. This is represented by a significant decrease in AP fall time (Fig. 2D) and a decrease in the AP half-width (Fig. 2E). This effect partially recovered by 24h. The AP shoulder has been attributed to calcium and sodium currents, potentially, due to the inability of these channels to completely inactivate during the AP falling phase^37^. There was no change in AP overshoot, amplitude or rise time (Table 1). Together, these data suggest that potential changes in voltage-gated ion channels (such as an increase in potassium channels and/or decrease in voltage-gated calcium and sodium channels) are modulating the generation and properties of APs to promote a decrease in neuronal excitability. The increase in rheobase and a decrease in the percentage of responding DRG neurons in ramp protocol but not in step protocol also suggests that adaptive changes may involve both voltage- and time-dependent gating processes (i.e. a slowly activating current).

**Figure 2:**
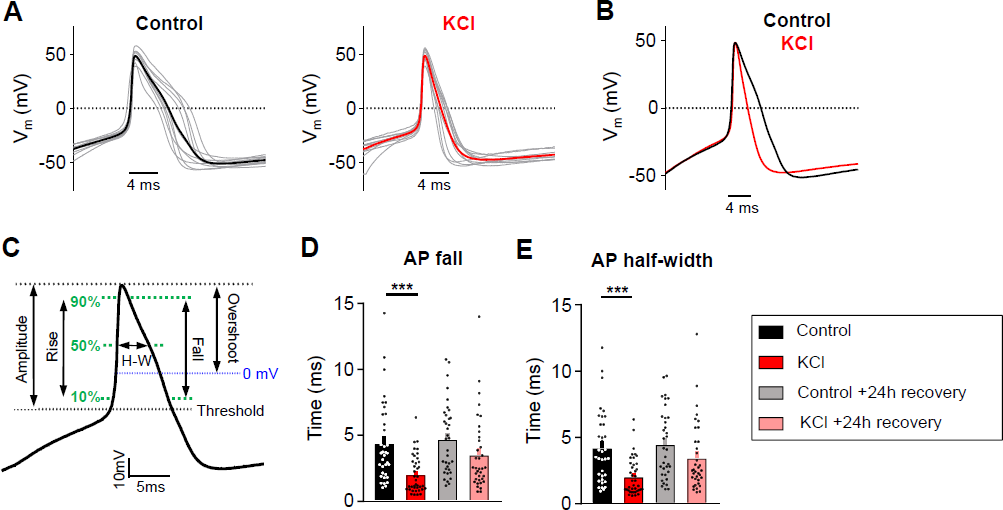
AP duration is altered in mouse DRG neurons following sustained depolarization. A) Current clamp AP traces from cultured mouse DRG neurons treated for 24hrs with control media (left) or 30mM KCl (right). Ten representative traces from each condition (grey) and average (bold color) are overlaid (aligned to peak depolarization) to display AP waveforms. The first AP at rheobase to step pulses was used. B) Average AP traces are overlaid to display difference with treatment (control=black, KCl=red). C) Diagram of AP waveform characteristics analyzed on an average AP trace. Analysis of AP fall (D) and half-width (E) from all mouse sensory neurons. One-way ANOVA with Tukey’s multiple comparisons. *p<0.05, **p<0.01, ***p<0.001, **** p<0.0001. Data are represented as mean ± SEM

**Table 1:** Impact of sustained depolarization on additional passive and active electrophysiological properties in mouse sensory neurons, related to figure 1 and 2.

|  | Capacitance (pf) | V<sub>rest</sub> (mV) | AP Amplitude (mV) | AP Overshoot (mV) | AP Rise (ms) |
| --- | --- | --- | --- | --- | --- |
| **Control (n=36)** | 25.2 ± 2.2 | -57.5 ± 1.6 | 63.2 ± 2.3 | 47.5 ± 1.3 | 0.76 ± 0.08 |
| **KCl (n=39)** | 20.3 ± 1.2 | -57.8 ± 1.5 | 66.3 ± 1.6 | 47.5 ± 01.1 | 0.55 ± 0.04 |
| **Control +24h recovery (n=35)** | 22.5 ± 1.4 | -58.3 ± 1.4 | 66.2 ± 2.0 | 49.8 ± 1.2 | 0.73 ± 0.09 |
| **KCl +24h recovery (n=36)** | 22.7 ± 1.6 | -58.0 ± 1.6 | 62.8 ± 1.5 | 48.5 ± 1.2 | 0.68 ± 1.18 |

### Sustained depolarization alters the firing pattern of mouse DRG neurons

We identified three firing patterns in small DRG neurons: 1) “Single” spikers (only fire one AP at the start of current injection even with suprathreshold stimuli; also referred rapidly accommodating^38^), 2) “Delayed” spikers [do not fire APs for a least 100ms after the start of current injection (average of 578 ±73 ms)^38^, but lose their phenotypes at high suprathreshold stimuli (2-4x rheobase); also referred to as non-accommodating^38^], and 3) “Repeated” spikers (fire at the start of current injection and continue firing multiple APs during the course of current injection) (Fig. 3A). Analysis of the action potential waveform from these subtypes showed a significant increase in voltage threshold, a decrease in AP amplitude, and an increase in rise time in delayed spikers as compared to single and repeated spikers (Fig.3B**-C**). Single spikers also had a significantly increased threshold, decreased AP amplitude and decreased overshoot compared to repeated spikers (Fig. 3C). Regardless of treatment, the majority of cells with a single spiker phenotype to step depolarizations were NR cells (Fig. 1B,D) to ramp depolarizations (36 NRs/57 single spikers, or 63.2%); all of the NRs were single spikers (36 single spikers/36 NRs, or 100%). The proportion of the three firing subtypes was significantly altered in KCl–treated cells (Fig. 3D), such that there was a significant increase in the proportion of single spikers and a decrease in delayed and repeated spikers in KCl-treated cells compared to cells in control media (Fig. 3D). Delayed spikers were almost absent in KCl-treated cells (1/39 or 2%). These alterations in neuronal firing after sustained depolarization were recovered to control levels in cells placed in fresh media within 24h (Fig. 3D). AP waveforms were analyzed by subtype to determine whether the changes in AP characteristics with KCl treatment (loss of AP shoulder quantified as decreased input resistance, AP fall time and half-width) were due to the shift in firing patterns (i.e., an effect from just having more single firer AP waveforms) or whether KCl treatment impacted all subtype waveforms. Both single (Fig. 3E, F) and repeated (Fig. 3G, H) spikers showed a decrease in input resistance, AP fall time, and half-width with KCl treatment. Combined with the almost complete elimination of the delayed subtype after KCl (1/39 cells), this data suggests that the overall AP waveform changes with KCl treatment are due to alterations in all subtypes, in addition to the increase in the proportion of single spikers.

**Figure 3:**
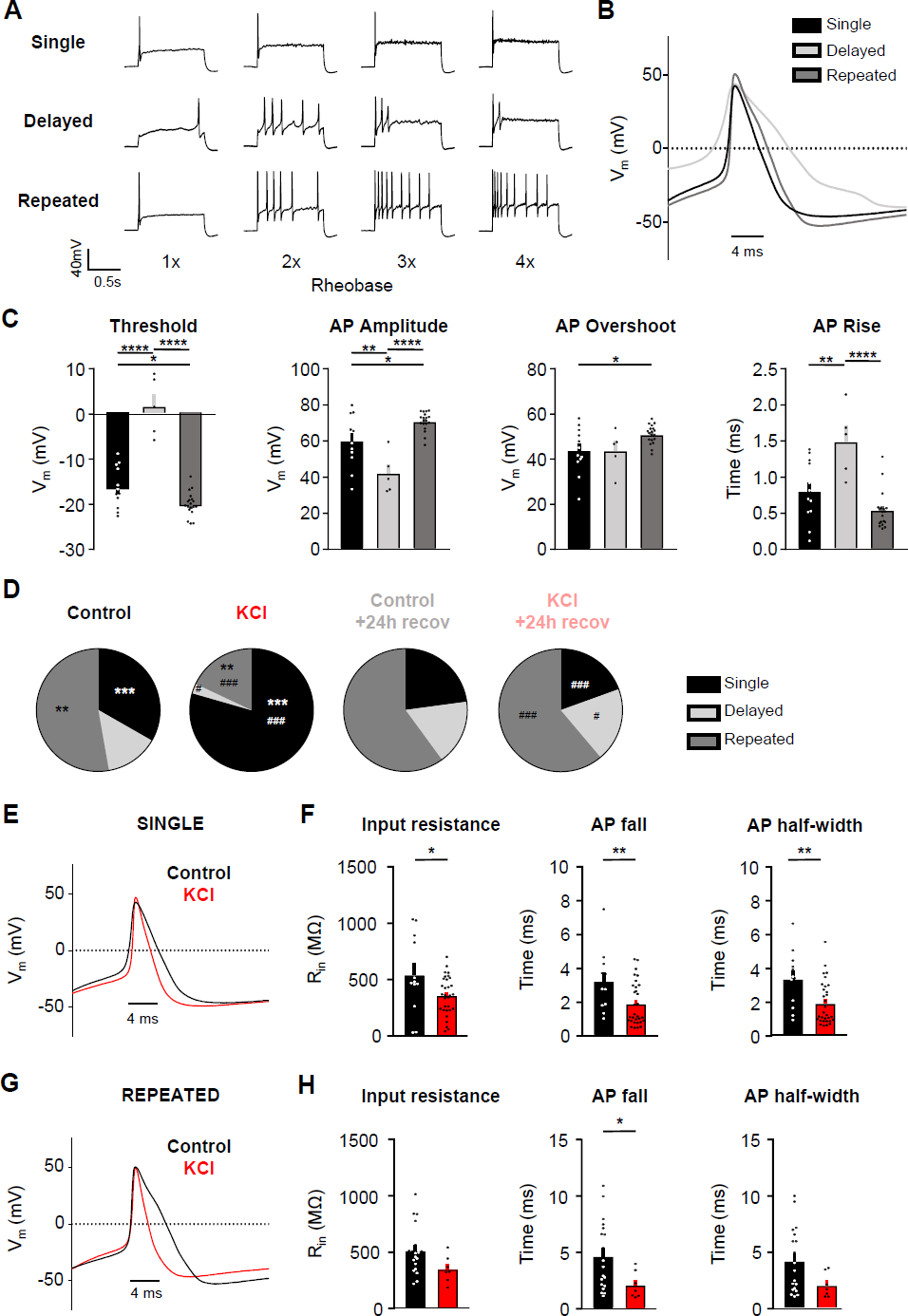
Sustained depolarization alters the firing pattern of mouse DRG neurons. A) Example current-clamp traces of single, delayed, and repeated subtypes of firing patterns in response to suprathreshold stimuli (1-4 times rheobase injections with step pulses). B) AP waveform traces (average of 5 representative traces per subtype; single=black, delayed=light grey, repeated=dark grey) overlaid to display differences between subtypes. The first AP at rheobase to step pulses was used. C) Analysis of threshold, AP amplitude, AP overshoot, and AP rise from all sensory neurons. All data were analyzed from the control treatment (single: n=12, black; delayed: n=5, light grey; repeated: n=19, dark grey). Statistical analysis was performed with one-way ANOVA with Tukey’s multiple comparisons. D) Relative proportions of firing pattern subtypes are displayed for each treatment group. Statistical analysis was performed with Chi-squared and Fisher’s exact t-tests. *=significant difference between control and KCl groups; ^#^=significant comparison between KCl and KCl+24h recovery groups. E) AP waveform traces (average of 5 representative traces per treatment; control=black, KCl=red) overlaid to display differences between treatments. The first AP at rheobase to step pulses was used. F) Analysis of input resistance, AP fall, and AP half-width of single firers from control (n=12, black) and KCl (n=31, red) treated groups. Statistical analysis was performed with student t-tests. G-H) Same analysis as E and F, but in repeated firers (control: n=19, KCl: n=7). There was only one delayed firer in the KCl-treated group, so AP analysis could not be performed for the delayed subtype. */^#^ p<0.05, **/^##^ p<0.01, ***/^###^ p<0.001, ****/^####^ p<0.0001. Data are represented as mean ± SEM

### Sustained activity did not alter neuronal excitability in mouse nociceptors

Exposure to 30mM KCl produced sustained depolarization, but not persistent AP firing (Fig. 1A, **bottom right**). However, the predominant phenotype seen in sensory neurons from rodents or humans with chronic pain is spontaneous firing and increased excitability, but minimal changes in membrane potential^14, 16, 25^. To test whether increased AP firing produces homeostatic changes, we used prolonged optogenetic stimulation of mouse sensory neurons (Fig. 4A). DRG neurons were cultured from transgenic mice expressing Channelrhodopsin-2 (ChR2) in TRPV1-lineage neurons (TRPV1:ChR2 mice; see Methods^39–41^). To generate sustained AP firing, DRG neurons were exposed to pulsed blue light (470nm illumination at 1Hz, 10ms pulses) for 24h in a tissue culture incubator, while control TRPV1:ChR2+ DRG neurons were kept in the dark. A 1 Hz stimulus frequency was selected because it is similar to the physiological firing frequency of spontaneously active nociceptors in rodent neuropathic pain models^42^. Additionally, stimulation at this frequency in hippocampal slices has been shown to induce homeostatic plasticity^43^ and produces nociceptive behaviors in TRPV1:ChR2 mice^44^. We confirmed that cells fire reliably to stimulation at 1Hz (7/8 with 100% fidelity), but not to higher frequencies (e.g. 10 Hz stimulation, 0/8 at 100% fidelity, Fig. 4B). After 24h of 1Hz stimulation, blue light illumination still evoked peak currents similar to those observed in control cells kept in the dark (Fig. 4C), suggesting there was no desensitization or bleaching of ChR2 during this prolonged period of stimulation.

**Figure 4:**
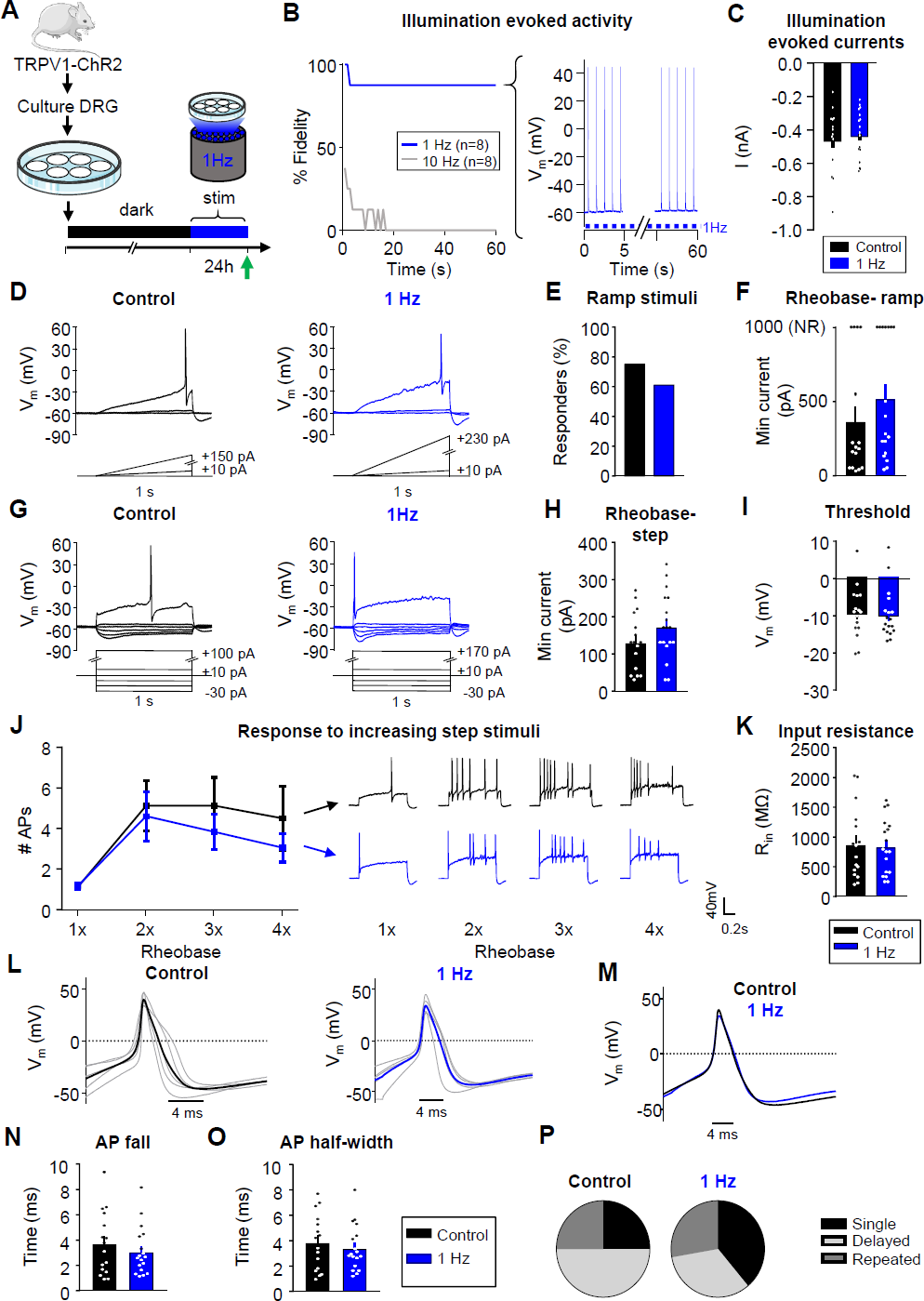
Sustained activity did not alter neuronal excitability in mouse nociceptors. A) Experimental design for testing homeostatic regulation of intrinsic excitability to sustained activity in cultured small DRG neurons from TRPV1:ChR2 transgenic mice. B) Left: Survival curve of nociceptors exposed to 470nm blue illumination at 1 or 10Hz, 10ms pulses. At 1min, 7/8 cells were still reliably firing with 1Hz stimulation. Right: example trace of a cell firing at 1Hz. C) A 1sec pulse of continuous illumination evoked similar peak currents in control cells kept in the dark (n=16; black) and cells stimulated with 24hrs illumination at 1Hz on an LED array (n=18; blue). Excitability was analyzed as described in Fig 1, but with treatment groups to test a sustained activity stimulus. Comparison of example current clamp traces (D), the proportion of responders (E), and rheobase (F) determined from ramp current injections. Comparison of example current-clamp traces (G), rheobase (H), threshold (I), response to suprathreshold stimuli (J), and input resistance (K) determined from step current injections. Treatment groups: cells kept in the dark (n=16; black), cells exposed to 24h illumination at 1Hz (n=18; blue). All cells were cultured from TRPV1:ChR2 mice. L) Current clamp AP traces from nociceptors treated for 24hrs in the dark (left) or 1Hz stimulation (right). Eight representative action potential traces from each condition (grey) and average (bold color) are overlaid. The first AP at rheobase to step pulses was used. M) Average AP traces are overlaid to show similar AP waveforms. Analysis of AP fall (N) and half-width (O) from all nociceptors. P) Relative proportions of firing pattern subtypes are displayed for each treatment group. Data are represented as mean ± SEM

Excitability data were collected from 34 small diameter (19 ± 0.2μm), ChR2-positive DRG neurons from 6 (3 male, 3 female) mice. In contrast to sustained depolarization, sustained 1Hz optogenetic activation did not significantly change the number of responders or affect rheobase during ramp protocol (Fig. 4D-F). In addition, there were also no significant differences in rheobase, threshold or response to suprathreshold stimuli to the step protocol (Fig. 4G-J), thus confirming that sustained activity did not alter intrinsic excitability. We also did not find changes in input resistance (Fig. 4K), AP fall or half width (Fig. 4L-O) nor in the proportion of firing pattern subtypes in DRG neurons (Fig. 4P). There were also no changes in other AP properties or passive membrane properties (Table 2).

**Table 2:**
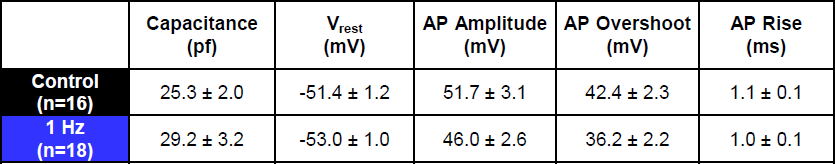
Impact of sustained activity on additional passive and active electrophysiological properties in mouse nociceptors, related to figure 4.

|  | Capacitance (pf) | V<sub>rest</sub> (mV) | AP Amplitude (mV) | AP Overshoot (mV) | AP Rise (ms) |
| --- | --- | --- | --- | --- | --- |
| **Control (n=16)** | 25.3 ± 2.0 | -51.4 ± 1.2 | 51.7 ± 3.1 | 42.4 ± 2.3 | 1.1 ± 0.1 |
| **1 Hz (n=18)** | 29.2 ± 3.2 | -53.0 ± 1.0 | 46.0 ± 2.6 | 36.2 ± 2.2 | 1.0 ± 0.1 |

It is interesting to note that even under control conditions, there were significant differences (p<0.05, Chi-squared) in the proportions of firing patterns between C57Bl6 (Fig.3D “media”) and TRPV1:ChR2 mice (Fig. 4P, “dark”). There were significantly more (p<0.05, Fisher’s exact) delayed spikers in neurons from TRPV1:ChR2 mice than C57Bl/6. It is possible that the TRPV1-Cre line biases the experimenter to a more homogenous DRG neuron population compared to our recordings from the C57Bl/6 mice.

### Sustained depolarization induces homeostatic plasticity in human DRG neurons

We next tested whether similar homeostatic mechanisms are engaged in human DRG neurons after sustained depolarization. Dorsal root ganglia were obtained from organ donors (Table 3). The experimental design was similar to that used for mice (Fig. 5A). At 3-5 DIV (after glia migrated off the cultured neuronal membranes^45^), human DRG neurons were incubated in 30mM KCl or control media for 24h. In contrast to mouse DRG neurons, 5/6 or 83% of human DRG neurons fired a series of APs during the rapid onset of depolarization in response to 30mM KCl, but we did not observe sustained AP firing during the course of KCl treatment (Fig. 5B). After this transient firing, we observed sustained depolarization to −6 ± 1mV (n=6) in human DRG neurons (Fig. 5B, **inset**).

**Table 1:**
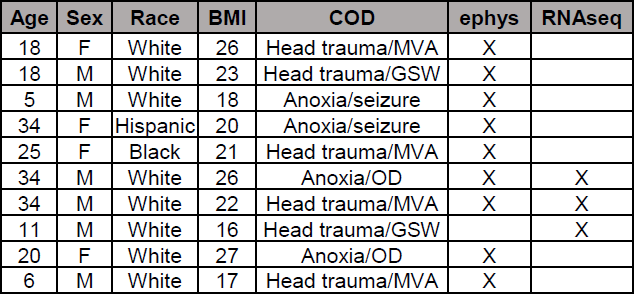
Human DRG donor characteristics, related to figures 5 and S2.

**Figure 5:**
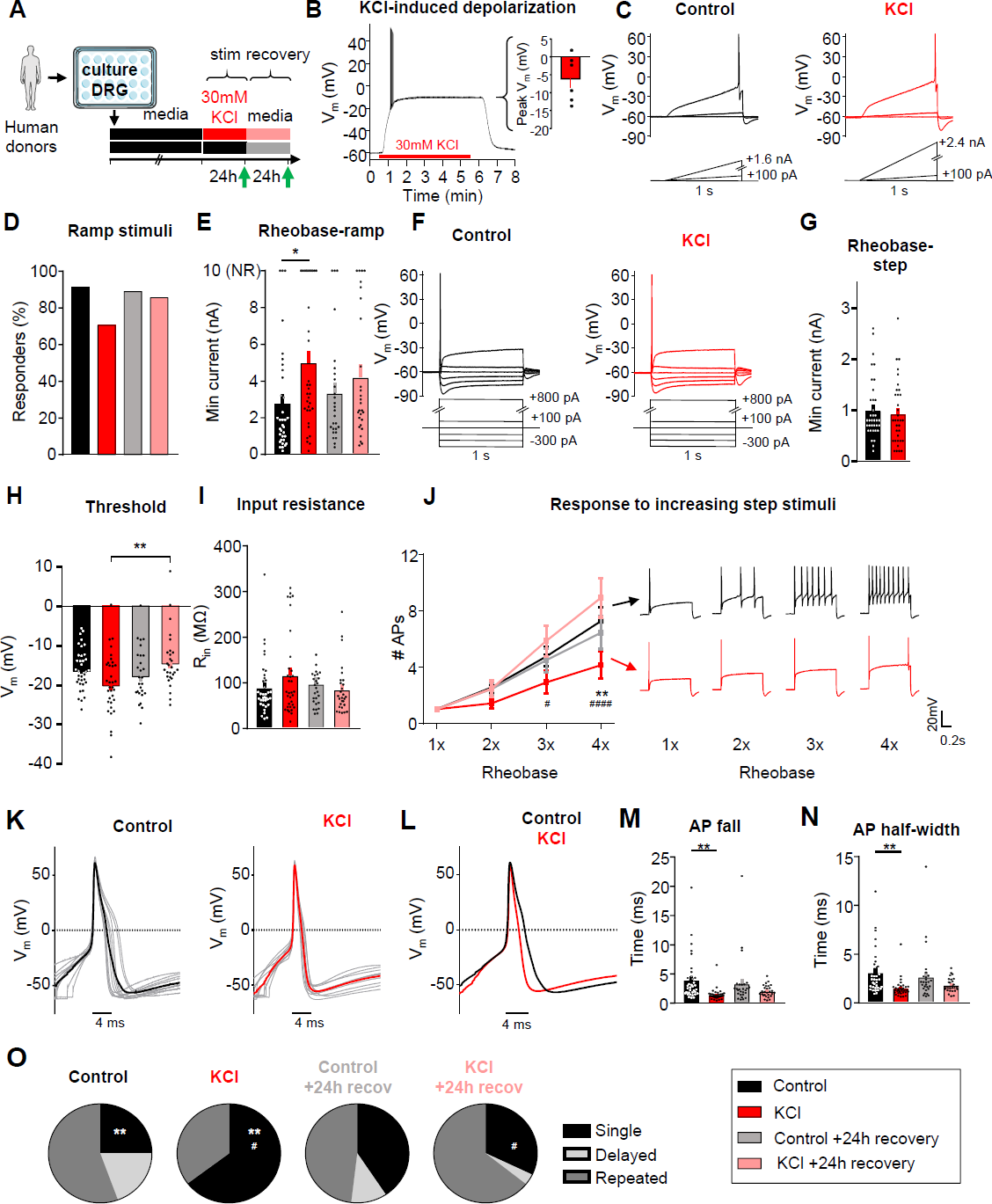
Sustained depolarization induces homeostatic plasticity in human DRG neurons. Experimental design for testing homeostatic regulation of intrinsic excitability to a sustained depolarization stimulus in cultured, small/medium diameter DRG neurons from human tissue/organ donors (A). B) Left: example current-clamp trace from acute, 5min application of 30mM KCl to a human nociceptor and subsequent washout of effect. Right: KCl application resulted in a sustained depolarization to −6 ± 1mV (n=6). Excitability was analyzed as described in Fig 1, but with data collected from human sensory neurons. Comparison of example current clamp traces (C), the proportion of responders (D), and rheobase (E) determined from ramp current injections. Comparison of example current-clamp traces (F), rheobase (G), threshold (H), input resistance (I), and response to suprathreshold stimuli (J) determined from step current injections. Treatment groups: 24h in control, media alone (n=36; black), 24h in media supplemented with 30mM KCl (n=34; red), 24h control followed by an additional 24h in fresh media (n=27; grey), and 24h 30mM KCl followed by an additional 24h “recovery” in fresh media (n=28; pink). K) Current clamp AP traces from cultured mouse DRG neurons treated for 24hrs with control media (left) or 30mM KCl (right). Ten representative traces from each condition (grey) and average (bold color) are overlaid to display AP waveforms. The first AP at rheobase to step pulses was used. L) Average AP traces are overlaid to display differences with treatment (control=black, KCl=red). Analysis of AP fall (M) and half-width (N) from all DRG neurons. O) Relative proportions of firing pattern subtypes are displayed for each treatment group. *=significant difference between control and KCl groups; ^#^=significant comparison between KCl and KCl+24h recovery groups; one-way and two-way ANOVAs with Tukey’s multiple comparisons. Chi-squared and Fisher’s exact tests were also performed. */^#^ p<0.05, **/^##^ p<0.01, ***/^###^ p<0.001, ****/^####^ p<0.0001. Data are represented as mean ± SEM

Data were collected from 125 small diameter (42 ± 1μm) DRG neurons from 9 (5 male, 4 female) human donors (Table 3). Excitability was assessed in human DRG neurons (small to medium diameter, <60 μm^46–48^) after 24 ± 4hrs of incubation in 30mM KCl in media, or after a further 24 ± 4hrs recovery in fresh media (Fig. 5A, **green arrows**). Using ramp current injections (Fig. 5C), we found that the percentage of KCl-treated human DRG neurons responding to current injection was not statistically significant from control media (Fig. 5D, p=0.09). However, similar to mouse, KCl-treated human DRG neurons had a significantly increased rheobase to ramp depolarizations compared to control media, which was partially recovered within 24h (Fig. 5E). Non-responder (NR) DRG neurons did not fire an AP up to the maximum ramp current of 10 nA (Fig 5E**-top**). While only a proportion of neurons responded to ramp protocol, all neurons responded to step current injections (Fig. 5F). There were no significant changes in rheobase, AP threshold and input resistance (Fig. 5G-I), but KCl-treated human DRG neurons showed a significant decrease in the number of APs fired in response to increasing step stimuli at 4x rheobase. This change recovered after 24h in fresh control media (Fig. 5J). Therefore, the increase in rheobase-ramp and the decrease in AP firing to suprathreshold step stimuli demonstrate an overall decrease in excitability of human DRG neurons following a sustained depolarizing stimulus, similar to mouse DRG neurons.

There were no significant changes with KCl treatment on the passive membrane properties of human DRG neurons (Table 4). However, as with mouse DRG neurons, analysis of AP characteristics revealed a significant decrease in AP fall time and half-width in KCl-treated cells, which partially recovered within 24h (Fig. 5K-N, Table 4). We also identified three firing patterns in human DRG neurons similar to mouse DRG neurons (**Fig.S1**). Analysis of the AP waveform from these subtypes showed a significant increase in voltage threshold in delayed spikers compared with single spikers and repeated spikers and a significant increase in AP rise time of delayed spikers compared with repeated spikers (Fig. S1). We did not observe significant changes in AP amplitude and AP overshoot. Additionally, there was a significant increase in the number of single spikers in KCl-treated cells, which also recovered within 24h (Fig.5O). Together, this data show that a sustained depolarizing stimulus is also capable of triggering intrinsic homeostatic plasticity in human sensory neurons.

**Table 4:** Impact of sustained depolarization on additional passive and active electrophysiological properties of human DRG neurons, related to figure 5. AP overshoot was statistically significant when comparing KCl group versus Control + 24hrs recovery and KCl group versus KCl + 24hrs recovery. ^#^p&lt;0.05, one-way ANOVA with Tukey’s multiple comparisons. Data are represented as mean ± SEM

|  | Capacitance (pf) | RMP (mV) | AP amplitude (mV) | AP Overshoot (mV) | AP rise (ms) |
| --- | --- | --- | --- | --- | --- |
| **Control (n=36)** | 123 ± 11 | -58 ± 1 | 76 ± 1 | 60 ± 1 | 0.53 ± 0.04 |
| **KCl (n=34)** | 118 ± 13 | -60 ± 1 | 72 ± 3 | 52 ± 2<sup>#</sup> | 0.52 ± 0.05 |
| **Control +24h recovery (n=27)** | 120 ± 8 | -60 ± 1 | 77 ± 1 | 59 ± 1 | 0.47 ± 0.04 |
| **KCl +24h recovery (n=28)** | 131 ± 10 | -58 ± 1 | 73 ± 2 | 59 ± 2<sup>#</sup> | 0.49 ± 0.03 |

### Assessment of global gene expression in mouse and human DRG neurons following sustained depolarization

As an unbiased approach to identify potential contributors to this homeostatic plasticity, we performed bulk RNAseq on mouse and human DRG cultures (n=3 males, each) to assess global gene expression changes after 24h treatment with 30mM KCl (or control media), and 24h recovery in fresh media (Table 3). A total of 21,962 genes (14,253 expressed) were analyzed in mouse, and 20,344 genes (14,656 expressed) in human (see methods^49–52^) . Hierarchical clustering showed that there were negligible alterations between treatment conditions in the global transcriptome across mouse and human DRG neurons (Fig. S2A-B). In fact, the individual subject variance was greater than the variance caused by the treatment, with the lowest correlation still greater than 0.86 (mouse) and 0.92 (human). Individual gene analysis showed no significant changes with treatment for human (Fig. S2C) or mouse (Fig. S2D) DRG neurons. A closer analysis of individual potassium and sodium channels did not reveal significant changes between treatments as well (Fig. S2C-D, **middle and right**).

### Sustained depolarization of mouse DRG neurons reduces voltage-gated sodium currents

Because no clear changes were observed in our bulk RNAseq analysis, we took a candidate-based approach to identify potential mechanisms of the observed homeostatic plasticity. Voltage-gated sodium and potassium channels are the key contributors to AP generation and waveforms ^53^, and thus represent the most likely mediators of the changes in intrinsic excitability of mouse and human sensory neurons. Mouse DRG neurons were incubated in 30mM KCl for 24h as shown in Figure 1A. After 24h, we performed voltage clamp recordings to assess the impact of sustained depolarization on voltage-gated sodium and potassium currents.

Total voltage-gated sodium currents were significantly reduced in cells treated with KCl compared to control media when analyzed using a voltage ramp protocol^54^ (Fig. 6A-B), or a step-pulse protocol (Fig. 6C-E). The notably larger reduction in sodium currents measured in the ramp protocol compared to the step protocol suggests that a time-dependent property in sodium channel gating might contribute to the reduction in excitability. This is consistent with the data shown in Fig. 1B and 1E, where ramp depolarization revealed more substantial differences after prolonged KCl treatment compared to step depolarization in the current clamp experiments.

**Figure 6:**
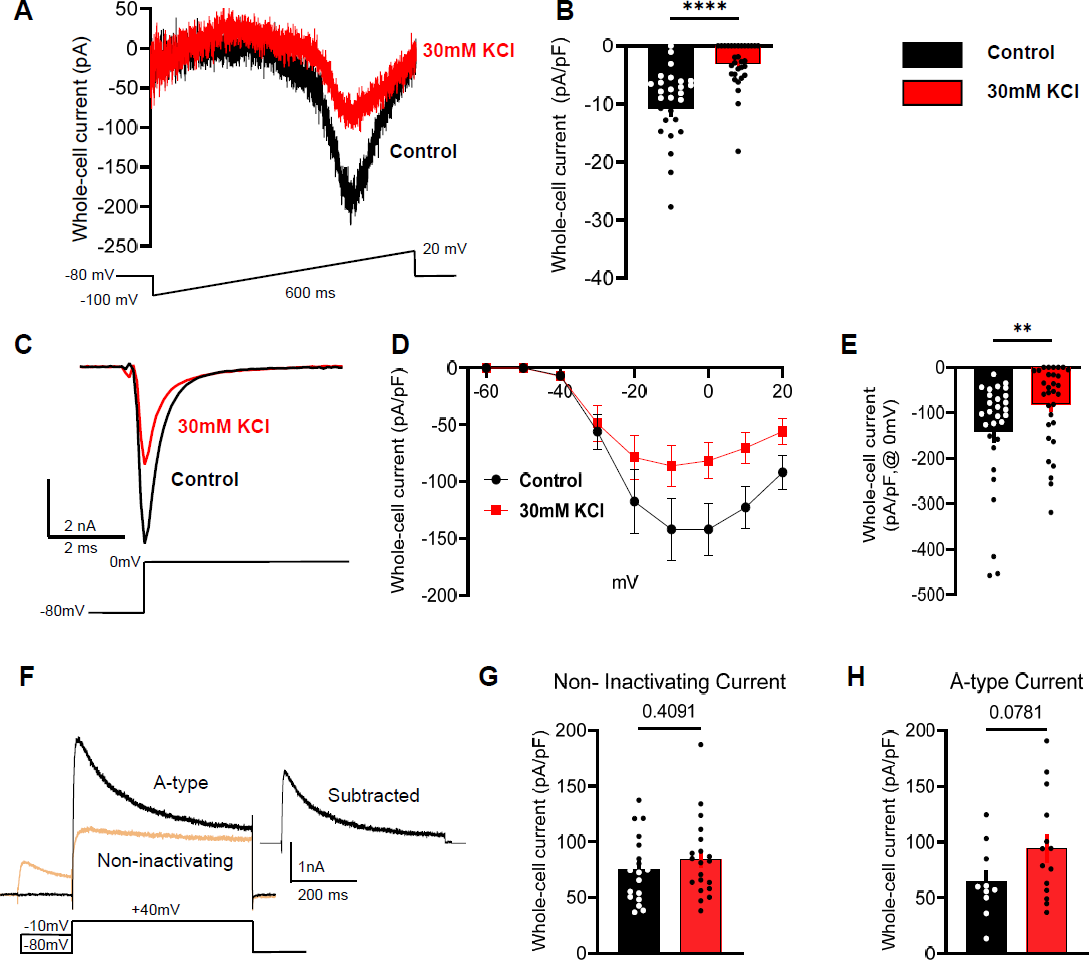
Sustained depolarization of mouse DRG neurons inhibits voltage-gated sodium channel currents. A) Example traces of voltage-gated sodium currents from control-treated cells (black) and KCl-treated cells (red) using a voltage-ramp protocol shown below traces. Cells were held at −80 mv, and ramped from −100 to +20 mV for 600 ms. B) Statistical analysis of control cells (black, n=28) versus KCl-treated cells (red, n=31). C) Example traces of voltage-gated sodium currents from control-treated cells using a voltage-step pulse protocol shown below traces. D) I/V curve of voltage-gated sodium currents from control and KCl-treated cells. Cells were held −80 mV and a series of step pulses ranging from −60 to +20 mV were applied in +10 mV increments for 1s. E) Current density analysis of control cells (black, n=29) versus KCl-treated cells (red, n=31) at 0mV. F) Example traces of A-type and non-inactivating voltage-gated potassium currents in mouse DRG neurons using a voltage-step pulse protocol shown below traces. Cells were clamped at −70 mV and conditioning prepulses ranging from −120 to +40 mV were applied in +10 mV increments for 150 ms every 5s followed by a pulse step of +40 mV for 500 ms. The isolation protocol of A-type voltage-gated potassium currents is shown below the representative traces. G-H) Current density analysis of A-type currents and non-inactivating currents (control-black, KCl–red). Statistical significance was calculated with the Mann-Whitney test or unpaired t-test. Scatter plots and mean ± SEM for current amplitudes.*p<0.05, **p<0.01, ***p<0.001, ****p<0.0001. Data are represented as mean ± SEM

We also observed an unidentified outward current that was slightly increased in KCl-treated cells during the isolation of sodium currents in the step-pulse protocol. This unblocked outward current cannot be attributed to potassium since potassium ions were omitted from the intra and extracellular solutions (Fig. S3).

We further investigated if there were changes in current density from the two main types of potassium currents present in DRG neurons: a fast inactivating (A-type) and a non-inactivating component^55, 56^. We isolated both currents and compared the peak and steady current density in KCl and control-media neurons. There was no difference in the non-inactivating component of whole-cell potassium currents, and while there was a trend toward an increase in A-type current density in the KCl-treated neurons, this was not statistically significant (Fig. 6F-H).

## DISCUSSION

The cellular and molecular mechanisms that dictate Hebbian and homeostatic plasticity have been widely studied in the CNS due to their critical role in maintaining the efficiency of neuronal communication in the event of global perturbation of the overall synaptic network^1, 9, 57, 58^. However, whether similar modulatory mechanisms are present in peripheral sensory neurons has been largely unexplored. Interestingly, DRG neurons, unlike CNS neurons, do not form synapses *in vitro*, and this allowed us to differentiate effects of sustained depolarization versus activity using different stimuli (KCl vs optogenetic activation, respectively). It is important to note that both stimuli presented in these experiments were not designed to mimic physiological conditions during pain, but rather to stimulate peripheral sensory neurons and then determine if they could compensate by altering intrinsic excitability.

We found that small diameter mouse DRG neurons, representing putative nociceptors, showed significant inhibition of intrinsic excitability following prolonged depolarization, but not prolonged activity (Figs. 1,4**,5**). In our study, chronically depolarized mouse DRG neurons showed an increase in rheobase, decrease in input resistance and substantial decrease in action potential generation to suprathreshold current injections to up to four times rheobase compared to control-treated neurons. These findings are similar to reports of chronic depolarization of hippocampal and cortical neurons in culture using high concentration of KCl^2, 8^. However, we also found changes in AP profiles; sustained depolarization resulted in a decrease shoulder of the AP, measured as a decreased AP fall and half-width. The parameters of membrane excitability altered by sustained depolarization partially or fully returned to baseline within 24h. These findings show that mouse DRG neurons can exert homeostatic control to regulate their intrinsic excitability similar to neurons in the CNS.

One major hurdle to developing better treatments for chronic pain is improving translation of research findings in animals to successful clinical trials. A powerful approach is to determine if experimental findings using rodent neurons are mirrored in human neurons by examining neurons obtained from human organ donors^45^. Homeostatic plasticity was also engaged in human DRG neurons chronically depolarized with KCl (Fig. 5), suggesting that this homeostatic plasticity is conserved across species. Homeostatic plasticity has been observed in multiple species, including flies^59^, crustaceans^60, 61^ and rodents^2^. To our knowledge, our study is the first to provide evidence for homeostatic plasticity in primary human neurons. In addition, we found that the firing patterns of both mouse and human DRG neurons were significantly altered after sustained depolarization. Our results in mouse and human sensory neurons show three main types of firing patterns: single spikers (also referred to as rapidly accommodating), delayed spikers (also referred to as non-accommodating^38^) and repetitive spikers (Fig. 3,S1). Sustained depolarization shifted the firing pattern of both mouse and human sensory DRG neurons to preferentially single spikers (Fig. 3D and 5O), which correlates with inhibition of neuronal excitability as result of homeostatic plasticity.

Treatment with elevated concentrations of extracellular KCl resulted in sustained depolarization but not sustained activity of DRG neurons in culture, similar to observations from cultured CNS neurons^2, 8, 62^. We therefore studied whether similar homeostatic mechanisms are engaged in DRG neurons in response to cell autonomous increases in activity. Using an optogenetic approach, we photostimulated TRPV1-lineage nociceptors at a sustained rate of 1Hz for 24h and measured whether these DRG neurons showed changes in intrinsic excitability after prolonged firing, as shown previously in hippocampal pyramidal neurons^43^. We found that a sustained increase in AP firing was not sufficient to trigger homeostatic plasticity in sensory neurons (Fig. 4). While persistent firing at 1 Hz for 24h produces some degree of net depolarization, it is clear that this is insufficient to engage the homeostatic mechanisms observed with sustained depolarization in sensory neurons. This was surprising because pharmacological, electrical, and optogenetic stimuli have been used to study activity induced homeostatic plasticity in hippocampal and neocortical networks^2, 5, 6, 9, 10, 43, 63^. The absence of homeostatic plasticity in our sustained activity experiments could be due to inadequate simulation parameters in these experiments. First, it is possible that stimulation for more than 24 hours may be required. A previous study found that Complete Freund’s Adjuvant (CFA; an inflammatory stimulus) injected into the hindpaw of rodents decreased evoked AP firing frequencies of nociceptors at 8 weeks post injection but not at 2 days^64^, suggesting that >48hrs of activity may be required to induce plasticity in this model. Second, it is possible that a more physiological stimulus may be required, rather than artificially driving activity with optogenetics. DRG neurons fire in multiple patterns to different stimuli and may require a different pattern of input or activity to engage homeostatic plasticity mechanisms. Lastly, it is possible that mechanisms of intrinsic excitability were altered in this line of transgenic animals. Indeed, there was a difference in the proportions of neurons exhibiting different firing patterns between control C57Bl/6 and TRPV1:ChR2 mice. This suggests that the insertion of ChR2 into the membrane may alter excitability and firing patterns of nociceptors. This is an important observation that should be considered in future experiments studying nociceptors from these transgenic mice.

Our data show that human and mouse sensory neurons exert intrinsic homeostatic control in response to sustained depolarization. What mechanism of action is responsible for driving homeostatic plasticity in sensory neurons? Surprisingly, voltage clamp recordings showed that sustained depolarization of mouse DRG neurons elicit a profound decrease in voltage-gated sodium current density compared to control neurons (Fig.6). However, we did not observe any statistically significant changes in voltage-gated potassium current density, which further supports the critical role of sodium channels in the regulation of neuron excitability upon chronic depolarization. Interestingly, multiple reports showed that either sustained activation of sodium channels in cultured neurons or sustained elevation of intracellular calcium concentration in cultured adrenal chromaffin cells induces endocytosis and degradation of sodium channels, including destabilization of sodium channel mRNA^65, 66^. These processes involve activation of calpain, calcineurin and protein kinase C. Whether similar mechanisms are engaged in sensory neurons during homeostatic plasticity will demand additional experiments to define the underlying mechanisms of this reduction of sodium currents in sensory neurons produced by sustained depolarization.

We cannot eliminate the participation of other ion channels such as voltage-gated calcium channels, calcium-activated potassium channels, or various leak currents involved in the regulation of the process of homeostatic plasticity. Indeed, we noted a significant reduction in input resistance in mouse, but not human sensory neurons, and an increase in an unidentified outward current in neurons that underwent sustained depolarization compared to controls in our mouse voltage clamp experiments. These factors could contribute to the overall reduction in excitability and will require a more detailed investigation in future studies. However, the robust reduction of sodium currents is likely to contribute substantially to the overall inhibition of neuronal excitability in DRG neurons we observed here.

These data also expand our understanding of the extent to which different neuron types, from either the peripheral or central nervous system, utilize different intrinsic mechanisms to drive homeostatic plasticity. In cortical and hippocampal neurons, synaptic scaling of AMPA and NMDA receptors as well as changes in intrinsic excitability via voltage-gated calcium channels contribute to homeostatic mechanisms during periods of chronic disturbances in synaptic activity^1, 2, 43, 67^. Here, we show that DRG neurons appear to employ alterations in voltage-gated sodium channels to modulate intrinsic excitability during periods of sustained depolarization. DRG neurons express at least five different types of sodium channels (Na_v_s)^68, 69^ with Na_v_1.7, Na_v_1.8 and Na_v_1.9 representing the predominant subtypes^70^. In addition, Na_v_1.7, Na_v_1.8 and Na_v_1.9 are preferentially expressed in small diameter putative nociceptors, and are known to be linked to human pain syndromes ^68 71^. Na_v_1.7 is localized primarily in axons, where it promotes the initiation and conduction of action potential^70–72^. Na_v_1.8 and Na_v_1.9 show a more restricted expression in sensory neurons, where they produce the majority of the current responsible for the depolarizing phase of the APs (a critical step for the generation of multiple APs during ongoing neuronal activity) and the persistent TTX-resistant current found in small DRG neurons, respectively^37, 68, 73^. Our findings reveal that the total voltage-gated sodium current is reduced after chronic depolarization. If specific sodium channel subtypes are altered during homeostatic plasticity, these could represent important therapeutic targets for human chronic pain conditions.

Our RNAseq data revealed no significant changes in expression of individual potassium and sodium channel genes in KCl-treated groups compared to controls (Fig. S2). We can only conclude that at the RNA level, as detectable using bulk RNA sequencing, there are no significant changes in gene expression 24h after prolonged depolarization. We cannot exclude the possibility of altered trafficking, internalization, or post-translational modifications of voltage-gated sodium channels in producing the observed reductions in sodium current density. Moreover, it is important to note that RNAseq was performed on DRG cultures that contain glia and other non-neuronal cells. This is a potential confound that could mask changes in RNA expression in nociceptors. Single cell sequencing or traditional quantitative PCR approaches would be needed to specifically address this issue.

Our data support the concept that homeostatic plasticity is not restricted to neurons in the central nervous system, but it can also be engaged in peripheral somatosensory neurons. Whether these adaptive changes are engaged in the context of physiological or pathophysiological activation of sensory neurons is not known. Recent studies suggest that this may indeed be the case. In both rodent and human sensory neurons, prolonged exposure to inflammatory mediators has been shown to induce a similar suppression of neuronal excitability. For example, the pro-inflammatory cytokine macrophage migration inhibitory factor (MIF) depolarized and drove activity in putative nociceptors, and transitioned neurons from a hyperexcitable state (repetitive spiking phenotype) following acute stimulation, to a hypoexcitable state (single spiking phenotype) following prolonged exposure in rodents^74^. In addition, we recently found similar results in human DRG neurons in culture, where acute exposure to the inflammatory mediator bradykinin leads to hyperexcitability, while more prolonged exposure leads to hypoexcitability^75^.

Adaptive changes in firing patterns of peripheral sensory neurons do not happen in isolation, but rather are likely to impact overall function in the pain neuraxis. For example, *in vivo* recordings of DRG neurons using calcium imaging in models of chronic pain in mice suggest that sensory neurons synchronize their activity within hours after injury, and that this synchrony is necessary to drive cortical plasticity in somatosensory cortex^76^. Thus, adaptive changes in sensory neuron excitability appear to have implications for CNS circuit adaptations that are related to pain chronification.

These findings raise a number of questions regarding the relationship between peripheral homeostatic plasticity and the mechanisms responsible for driving the transition from acute to chronic pain. If nociceptors are capable of engaging homeostatic mechanisms that dampen intrinsic excitability, why are these mechanisms not engaged in the context of painful conditions associated with chronic nociceptor activity? Is effective homeostatic plasticity lost or disrupted during the transition to chronic pain, and further, can variability in the efficacy of homeostatic plasticity explain why some recover from injury, while others transition to chronic pain? These questions may be answered in future studies using animal models of chronic pain, and sensory neurons from human donors with a history of chronic pain.

## ACKNOWLEDGEMENTS

This study was supported by NIH grants DA007261 (L.A.M), NS113422 (J.S.D.R), NS065926 and U19NS130608 (T.J.P.), NS42595 and U19NS130607 (R.W.G. and B.A.C.). We thank Mid-America Transplant Services and John A. Lemen, as well as the donor families for making this research possible. We acknowledge the Genome Center and Yeunhee Kim, Ph.D. in UTD Research Core Facilities for mRNA library preparation and sequencing. We also thank Sherri Vogt for maintenance of animal colonies, and Judith Golden, Ph.D. for assistance with manuscript editing. Current affiliations: L.A.M.: Dept. of Neurobiology, University of Pittsburgh, Pittsburgh, PA and Hillman Cancer Center, University of Pittsburgh Medical Center, Pittsburgh, PA; T.D.S.: Pittsburgh Center for Pain Research, Department of Neurobiology, University of Pittsburgh; A.J.S.: Dept. of Symptom Research, Division of Internal Medicine, The University of Texas MD Anderson Cancer Center, Houston, TX 77030; A.W.: Department of Pediatrics, University of California, San Francisco.

## AUTHOR CONTRIBUTIONS

L.A.M., J.S.D.R., M.Y.P., B.A.C. and R.W.G. designed experiments. L.A.M., J.S.D.R., M.Y.P. and A.W. performed experiments and analyzed data. L.A.M., J.S.D.R., M.Y.P., T.D.S., A.J.S, R.A.S., J.A.L. and B.A.C collected mouse and human tissue. L.A.M., J.S.D.R. and R.W.G wrote the manuscript. L.A.M., J.S.D.R., M.Y.P., A.W., T.D.S., A.J.S, R.A.S. B.A.C., T.J.P, R.W.G. edited the manuscript.

## METHODS

### Animals

All procedures were approved by the Animal Care and Use Committee of Washington University and in strict accordance with the US *National Institute of Health (NIH) Guide for the Care and Use of Laboratory Animals*. Adult male and female mice (7-14 weeks of age) utilized in experiments were housed in Washington University School of Medicine animal facilities on a 12-hour light/dark cycle with access *ad libitum* to food and water.

For relevant optogenetic experiments, mice were generated with conditional expression of ChR2 in TrpV1 expressing sensory neurons by crossing heterozygous TrpV1-Cre mice (provided by Dr. Mark Hoon, NIDCR^39^) with homozygous Ai32 mice (Stock 3: 012569, The Jackson Laboratory)^40^. For the purposes of this study, we refer to these mice as “TRPV1:ChR2”. These mice were previously characterized in our lab^41^. All other experiments were performed using C57BL/6J mice bred in house, originally obtained from The Jackson Laboratory.

### Mouse DRG cultures

Mouse DRG cultures were performed as previously described^29, 30, 41^. Briefly, mice were deeply anesthetized with isoflurane and quickly decapitated or anesthetized with an i.p. injection of ketamine(100 mg/kg) and xylazine (10 mg/kg) cocktail prior to euthanasia. After confirming a lack of response to toe pinch, mice were perfused via the left ventricle with ice-cold Hank’s buffered salt solution. DRGs were removed from all spinal segments, and dissociated enzymatically with papain (0.33 mg/mL, Worthington) and collagenase type 2 (1.5 mg/mL, Worthington), and mechanically with trituration. Dissociated DRG were filtered (40 µM, Fisher) and cultured in DRG media [5% fetal bovine serum (Gibco) and 1% penicillin/streptomycin (Corning) in Neurobasal A medium 1x (Gibco) plus Glutamax (Life Technologies) and B27 (Gibco)] on glass cover slips coated with poly-D-lysine and collagen.

### Human DRG cultures

Human dorsal root ganglia extraction, dissection, and culturing was performed as described previously^45^. Briefly, in collaboration with Mid-America Transplant Services, L1-L5 DRG were extracted from tissue/organ donors less than 2 hrs after aortic cross clamp. Donor information is presented in Table 3. Human DRGs were placed in an *N*-methyl-D-glucamine (NMDG) solution for transport to lab, fine dissection, and mincing. Minced DRG were dissociated enzymatically with papain (Worthington) and collagenase type 2 (Sigma), and mechanically with trituration. Dissociated DRG were filtered (100µM, Fisher) and cultured in DRG media [5% fetal bovine serum (Gibco) and 1% penicillin/streptomycin (Corning) in Neurobasal A medium 1x (Gibco) plus Glutamax (Life Technologies) and B27 (Gibco)] on coated glass cover slips.

### Assessment of intrinsic homeostatic plasticity

At DIV4, mouse DRG cultures were treated for 24 ± 4hrs with a sustained depolarizing stimulus (30mM KCl, added to media) or sustained activity stimulus [470 nm illumination at 1Hz, 10ms pulses; culture plate placed on top of LED array in incubator (LIU470A, Thorlabs)]. Temperature of the media did not change after LED stimulation, indicating the LED array was not heating the media or cells. Coverslips were then changed to fresh media for 24 ± 4hrs recovery. For voltage clamp experiments, DRGs were tested after 24 ± 6hrs in vitro to improve space clamp. After each treatment timepoint, patch clamp electrophysiology was performed on cultured neurons to measure the response to sustained stimulation and recovery (Fig 1A, **4A**). Compensatory changes in excitability in response sustained depolarization or light stimulation were considered an indication of homeostatic regulation. Human DRG cultures were treated at DIV3-5, after glia had fallen off the neurons and exposed the cell membranes^6^. The investigator performing the physiology experiments was blinded to treatment condition.

### Whole-cell patch-clamp electrophysiology

Experiments were performed on small diameter, putatively nociceptive neurons (referred to as nociceptors) cultured from mouse (<30 μm^32–35^) and human (<60 μm^46–48^) DRG. Current clamp electrophysiological recordings were carried out in an external solution consisting of (in mM): 145 NaCl, 3 KCl, 2 CaCl_2_, 1.2 MgCl_2_, 7 Glucose and 10 HEPES, pH 7.3 with NaOH and ∼305 mOsm. Cells were recorded within 1 hour after being placed in external solution. As an exception, acute recordings demonstrating how cells respond in culture (Figs. 1C, 3B, and **4B**) were recorded in culture media as the external solution. Patch pipettes were pulled from thick-walled borosilicate glass (Sutter Instrument; Novato, CA) using a P-97 horizontal puller (Sutter Instrument), and had resistance values of 3-4 MΩ (mouse) and 2-3 MΩ (human). The intracellular solution contained (in mM): 120 potassium gluconate, 5 NaCl, 2 MgCl_2_, 0.1 CaCl_2_, 10 HEPES, 1.1 EGTA, 4 Na_2_ATP, 0.4 Na_2_GTP, 15 sodium phosphocreatine, adjusted to pH=7.3 with KOH, and 291 mOsm with sucrose. Fluorescence imaging and photostimulation at 470 nM was performed using a LED (M470L2, Thorlabs) coupled to the epi-fluorescence port of an upright microscope (BX50W1), Olympus). The LED was positioned for Kӧhler illumination, and light intensity was calibrated using a photodiode (S120C, Thorlabs) and power meter (PM100D, Thorlabs). Recordings were made using Patchmaster software (Heka Instruments; Bellmore, NY) controlling an EPC10 amplifier (Heka). All recordings were performed at room temperature with continuous, gravity-fed bath perfusion at a flow rate of 1-2mL/min. Data were sampled at 50kHz and analyzed off-line.

Changes in excitability were assessed in current clamp mode by 3 measures: rheobase, response to suprathreshold rheobase stimuli (accommodation), and action potential (AP) threshold. These were determined from a holding potential of −60 mV with a series of 1s depolarizing step (square pulse) or ramp current injections in increments of 10 pA (mouse, cut off at 1 nA) or 100 pA (human, cut off at 10 nA). Rheobase was defined as the minimum amount of current required to evoke a single AP. Response to suprathreshold stimuli was determined by counting the number of APs evoked at 1-4 times rheobase. Action potential threshold was defined as the mV/ms rate of change of the maximum depolarization reached before AP generation. Recordings were performed with a current clamp gain of 1pA/mV (mouse) or 10pA/mV (human).

The passive electrophysiological properties assessed were capacitance, input resistance (R_in_), and resting membrane potential (V_rest_). Capacitance was determined in voltage clamp mode using the compensation of the slow capacitive transient with the amplifier circuitry. Input resistance was determined in current clamp with a hyperpolarizing current injection of 10pA (mouse) or 100pA (human). V_rest_ was determined in current clamp as the average potential during a 1s pulse with no current injection. Neurons that exhibited a V_rest_ >-40mV or <-75mV were not analyzed. The active electrophysiological properties assessed were characteristics of the AP waveform. This included AP amplitude, overshoot, rise-time, fall-time, and half-width. The first AP evoked at rheobase from a step pulse was analyzed for these characteristics. AP overshoot was measured from 0mV to the peak depolarization, and AP amplitude was measured from threshold to peak depolarization. AP rise was measured as the time from 10-90% of the rising phase, AP fall was measured as the time from 10-90% of the falling phase, and AP half-width was measured as the time from 50% of the rising phase to 50% of the falling phase (Fig. 2C).

Voltage clamp electrophysiological recordings to isolate potassium currents were carried out in an external solution consisting of the following components (in mM) : 137 NaCl, 5 KCl, 1 MgCl2, 2 CaCl2, 10 HEPES, 10 Glucose, 2.5 CoCl2, 1 Lidocaine, 0.001 TTX. The intracellular solution contained (in mM): 140 K-gluconate, 10 HEPES, 5 EGTA, 1 MgCl2, 2 Na2 ATP. To isolate sodium currents, the external solution consisted of (in mM): 105 NaCl, 40 TEA-Cl, 10 HEPES, 13 Glucose, 1 MgCl2, 0.1 CdCl2. The intracellular solution contained (in mM): 110 CsCl, 10 EGTA, 0.1 CaCl2, 4 MgATP, 0.4 Na2GTP, 10 Na2Phosphocreatine, 10 HEPES, 10 TEA-Cl. For potassium current isolation, we voltage clamped the neurons at −70 mV and conditioning prepulses ranging from - 120 to +40 mV were applied in +10 mV increments for 150 ms every 5s^55^. This was followed by a voltage pulse step of +40 mV for 500 ms. Isolation of A-type currents was performed as shown in Fig.6. Moreover, for sodium current recording, mouse DRG neurons were clamped at −80 mV and a series of step pulses ranging from −60 to +20 mV were applied in +10 mV increments for 1s (step protocol). For ramp protocol of sodium current recording, mouse DRG neurons were held at −80 mv, and ramped from −100 to 20 mV for 600 ms as shown previously^54^.

### Pharmacological agents

All salts and pharmacological agents were mostly obtained from Sigma-Aldrich (St. Louis, MO) unless specified. Tetrodotoxin (TTX) citrate was obtained from Hello Bio (Princeton, NJ). A 3M stock of KCl, dissolved in DRG media with aliquots stored at −20°C until use, was diluted into culture DRG media for a final concentration of 30mM KCl.

### RNAseq

Mouse and human DRG cultured cells were scraped from 3 coverslips per treatment (media, KCl, KCl +recovery) into RNAlater and frozen at −80 °C until use. Samples were then thawed at room temperature, and cells pelleted at 5000 x g for 10 min. Cells were resuspended with 1 mL of QIAzol (QIAGEN Inc.; Germantown, MD) and transferred to tissue homogenizing CKMix tubes 2mL (Bertin Instruments; Montigny-le-Bretonneux, France). Homogenization was performed for 3 x 1 min at 20Hz at 4 °C. RNA extraction was performed with RNeasy Plus Universal Mini Kit from QIAGEN with the manufacturer provided protocol. RNA was eluted with 30 µL of RNase free water. Total RNAs were purified and subjected to TruSeq stranded mRNA library preparation according to the manufacturer’s instructions (Illumina; San Diego, CA). The libraries were quantified by Qubit (Invitrogen; Carlsbad, CA) and the average size of the libraries was determined using the High Sensitivity NGS fragment analysis kit on the Fragment Analyzer (Agilent Technologies; Santa Clara, CA). The normalized libraries were then sequenced on an Illumina NextSeq500 sequencing platform with 75-bp single-end reads for at least 20 million reads per sample in multiplexed sequencing experiments.

## QUANTIFICATION AND STATISTICAL ANALYSIS

### Whole-Cell Clamp Electrophysiology

Electrophysiological data were analyzed offline using custom-written macros and the Neuromatic plug-in^31^ in Igor Pro (WaveMetrics; Portland, OR) and also using Clampfit (Molecular Devices; San Jose, CA). Statistical analyses were performed with Prism software (GraphPad, La Jolla, CA) reported in figure legends. Changes in excitability and passive and active properties between groups and voltage clamp recordings were analyzed with one-way ANOVAs (Tukey multiple comparison), two-way repeated ANOVAs (Tukey multiple comparison), unpaired Student t-tests, Chi squared tests, or Fisher exact t-test, as appropriate. Data are presented as mean ± standard error of the mean (SEM).

### Bulk RNA-sequence

RNAseq Fastq files from the sequencing experiment were checked for quality by FastQC and trimming was done based on basepair-wise quality and per-base sequence content^49^. Trimmed Reads were then mapped against gencode mouse genome vM16 using STAR v2.2.1^50^.Transcripts per million (TPM) for every gene of every sample was quantified by stringtie v1.3.5 with output .bam files from STAR^51, 52^. Gene expression levels (TPM) were re-normalized against all protein coding genes to generate the new TPM for protein coding genes. A TPM <0.2 was below detection level and considered not expressed. Therefore, genes were only considered expressed when all 3 samples had TPMs>0.2 in at least one of the treatment groups (media, KCl, or KCl +24h recovery). Data were analyzed with paired student’s T-tests, with Benjamini-Hochberg multiple test correction. For graphs, the threshold of level of 0.2 TPM was added to all genes before taking the log transform.

---

**Figure S1:**
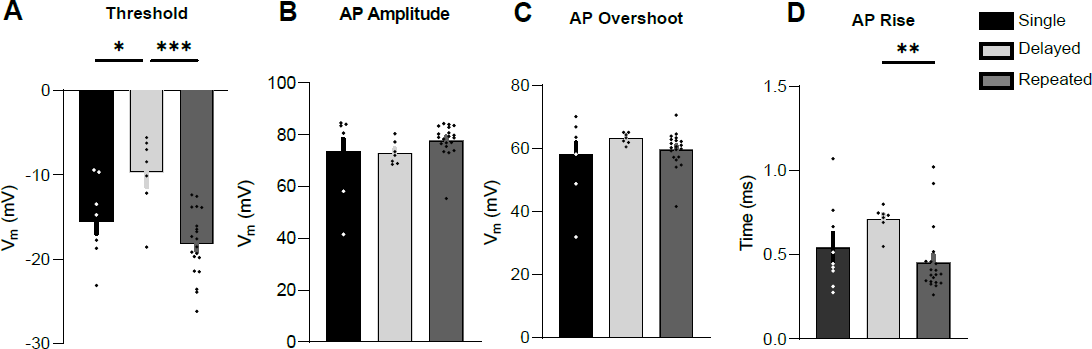
Analysis of active and passive properties of human DRG sensory neuron firing patterns, related to figure 5. A-D) Analysis of threshold, AP amplitude, AP overshoot, and AP rise from all human sensory neurons. All data were analyzed from the control treatment (single: n=9, black; delayed: n=7, light grey; repeated: n=20, dark grey). *p<0.05, **p<0.01, ***p<0.0001; one-way ANOVAs with Tukey’s multiple comparisons or Kruskal-Wallis test with Dunn’s multiple comparisons. Data are represented as mean ± SEM

**Figure S2:**
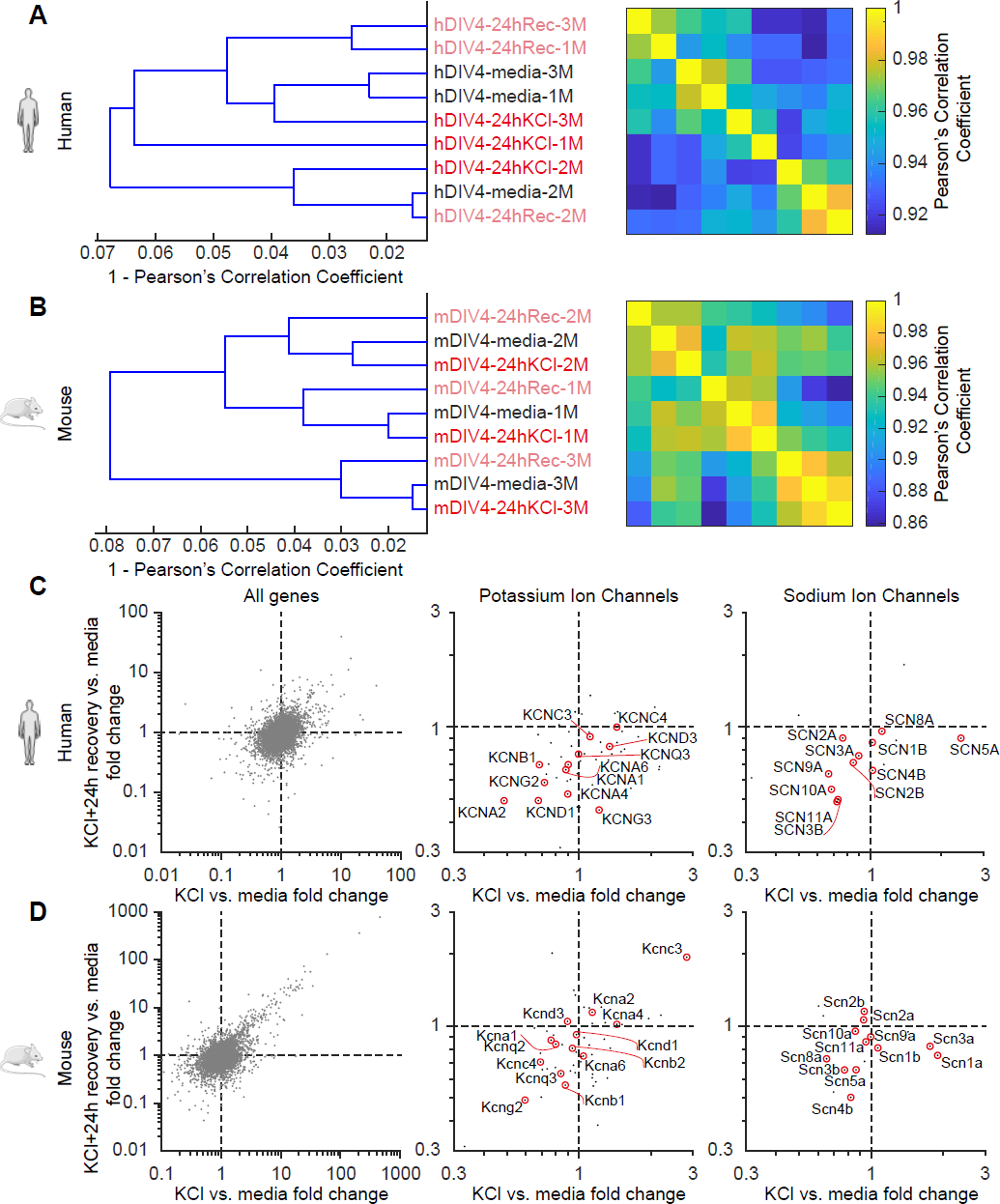
Assessment of global gene expression in mouse and human DRG neurons following sustained depolarization. A-B) Hierarchical clustering of RNA-seq samples from human and mouse DRG sensory neurons. N=3 cultures each, denoted by #1-3. Neurons were collected after approximately 4 days in vitro and the following treatment groups were studied: Media (control), 24hKCl (24h cultured in KCl media), and 24hRec (additional 24h recovery in fresh media after cultured in KCl for 24hr). C-D) Differential gene expression analysis between condition with KCl vs. media fold change (X-axis) and 24hRec vs media (Y-axis). Middle and right panels: Isolated potassium and sodium ion channels for better representation.

**Figure S3:**
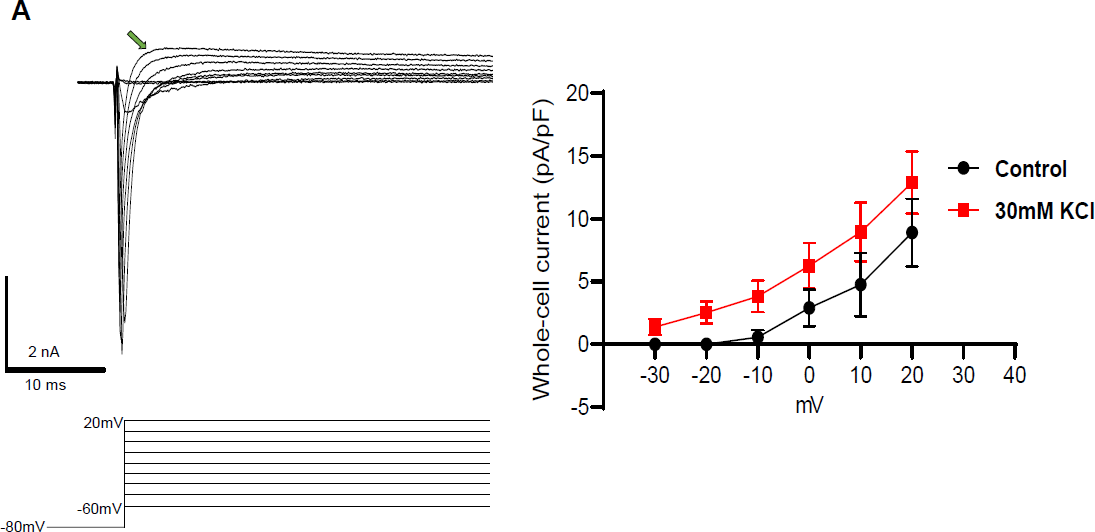
Mouse DRG neurons exhibit an unknown voltage-gated outward current, related to figure 6. A) Example traces of voltage-gated outward currents in mouse DRG neurons using a voltage-step pulse protocol shown below traces. Current density analysis of an unknown outward current from control (right; black) and KCl-treated cells (right; red). Analysis was done using the value at the peak of the outward current before the current decay (green arrow).

